## Supplemental Materials for "A preclinical model of THC edibles that produces high-dose cannabimimetic responses"

**Supplementary Table of Contents**

**Supplementary Figures**

Figure S1: CTR-gel and E-gel consumption by male and female mice

Figure S2: Analysis of THC-E-gel consumption behavior over 3-day access paradigm

Figure S3: Triad behavioral responses after CTR-gel and E-gel consumption

Figure S4: Triad behavioral responses to THC *i.p.* and E-gel administration by male and female mice

Figure S5: Methodology for THC-E-gel prediction of a behavioral response

**Supplementary Figure S1: CTR-gel and E-gel consumption by male and female mice**

**a**) Total E-gel consumed by males (blue) and females (orange) after 2 h *ad libitum* access. **b**) Calculated THC dose based on individual animal weights after E-gel consumption shown in a. No statistical significance due to Sex across doses found from Two-way ANOVA, Sidak’s post-hoc, N=2-11. Results are mean ± S.E.M.

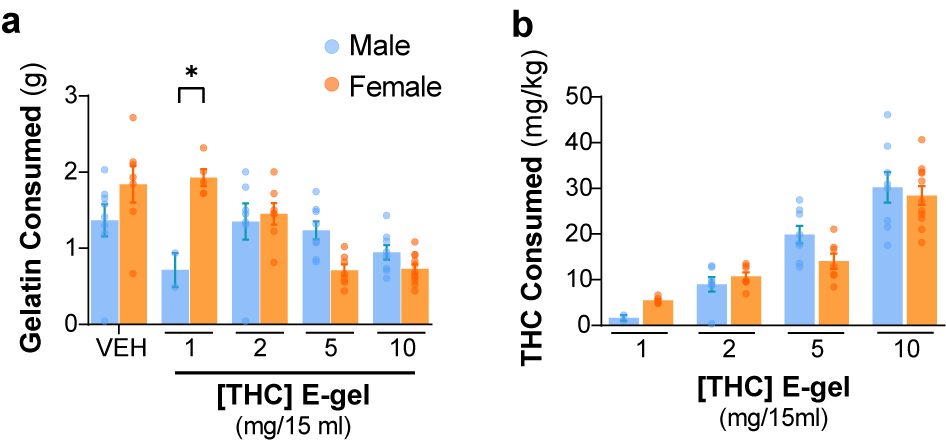

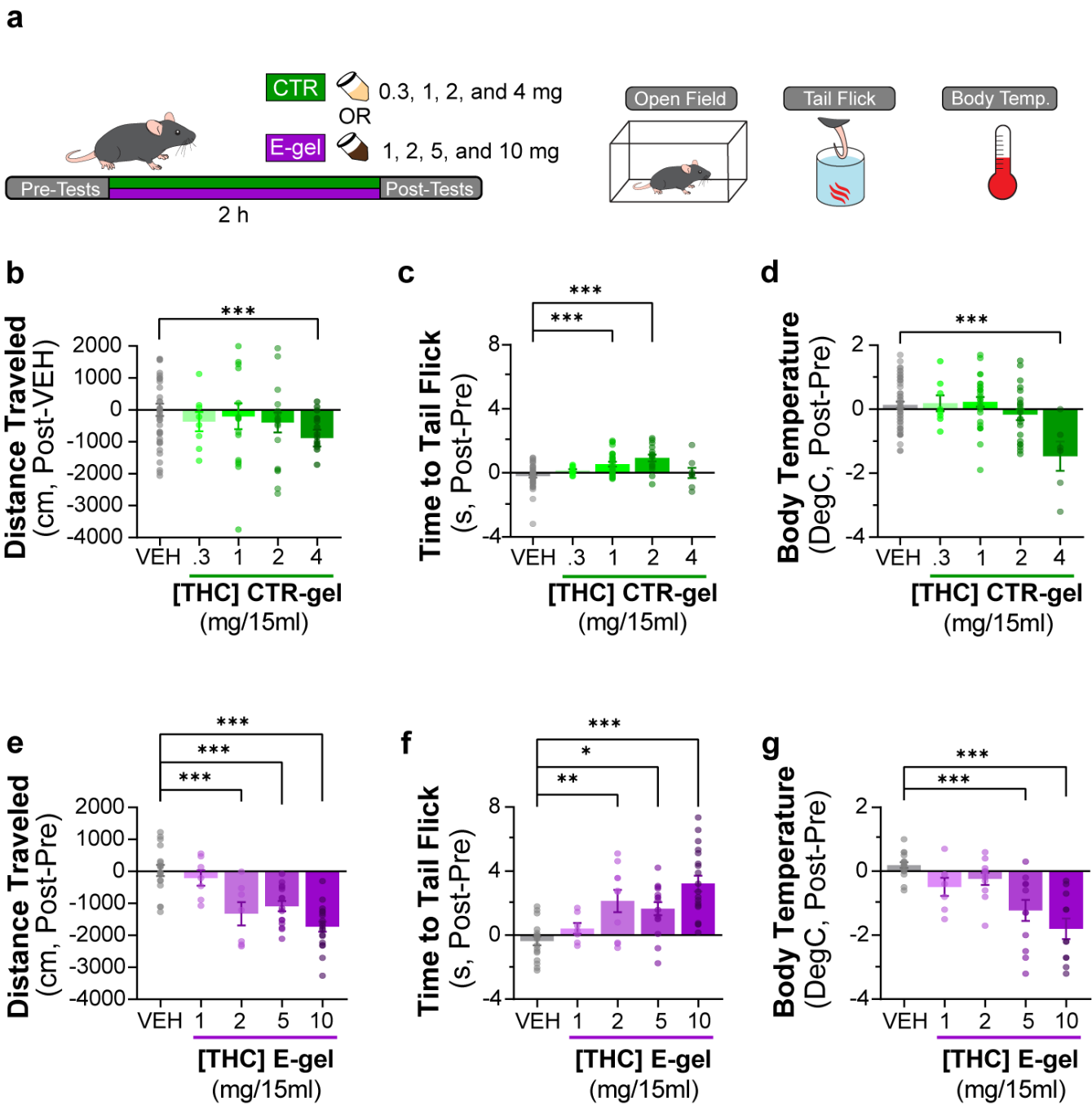

**Supplementary Figure S2: Triad behavioral responses after CTR-gel and E-gel consumption**

**a**) Diagram outlining 2 h exposure to either CTR-gel (green) or E-gel (purple) preceded and followed by Triad behavioral tests. **b-d**) Behavioral output immediately following CTR-gel administration for the Triad of cannabimimetic behaviors measuring hypolocomotion (**b**), analgesia (**c**), and hypothermia (**d**). In **b**, Distance traveled for CTR-gel was calculated based on averaged VEH consumption. All error expressed as SEM, unpaired Student’s T-test (*** p<0.001). **e-g**) Triad of cannabimimetic behaviors following THC E-gel administration measuring hypolocomotion (**e**), analgesia (**f**), and hypothermia (**g**). All error bars expressed as SEM, One-way ANOVA, Sidak’s post-hoc (* p<0.05, ** p<0.01, *** p<0.001).

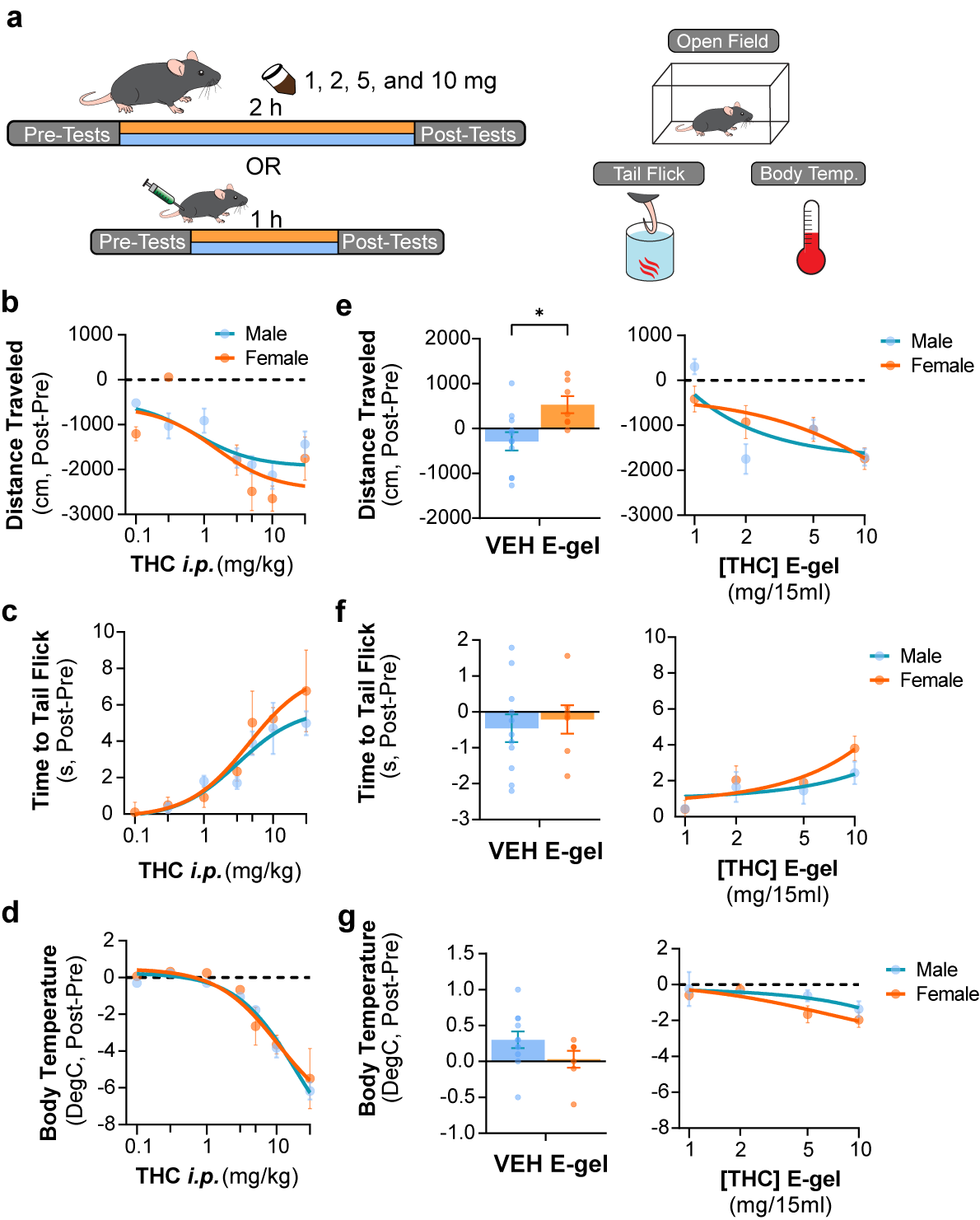

**Supplementary Figure S3: Triad behavioral responses to THC *i.p.* and E-gel administration by male and female mice**

**a**) Diagram outlining 2 h exposure to either E-gel or *i.p.* for males (cyan) or females (orange) preceded and followed by Triad behavioral tests. **b-d**) Dose-response curves for the Triad of hypolocomotion (**b**), analgesia (**c**), and hypothermia (**d**) cannabimimetic behaviors after *i.p.* THC administration with male responses shown in blue and female responses in orange. **e-g**) Male and female behavioral responses after access to THC E-gel doses for all three triad cannabimimetic behaviors: hypolocomotion (**e**), analgesia (**f**), and hypothermia (**g**). Sex-specific responses to VEH E-gel is separated from concentration-dependent curves. All error bars expressed as SEM, Two-way ANOVA, Sidak’s post-hoc, (* p<0.05).

**Supplementary Figure S4: Analysis of THC-E-gel consumption behavior over 3-day access paradigm**

**a**) Rate of consumption during the first 40 min of 10 mg/15ml THC E-gel access across the three-day paradigm where the THC group received vehicle E-gel on days 1 and 3. Error in S.E.M., unpaired Student’s T-test (* p<0.05). **b**) Rate of consumption after the first 40 min of E-gel access just as in a. Error in S.E.M., unpaired Student’s T-test (*p<0.05). **c**) Latency to start consuming gelatin across the same experimental paradigm in a showing a statistical significance between VEH and THC groups but not within experimental days, Two-way ANOVA, Sidak’s post-hoc, N=8 (* p<0.05).

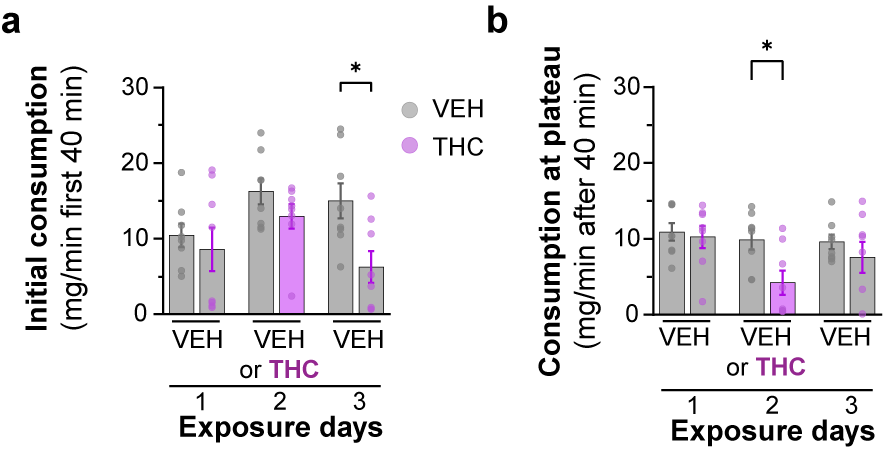

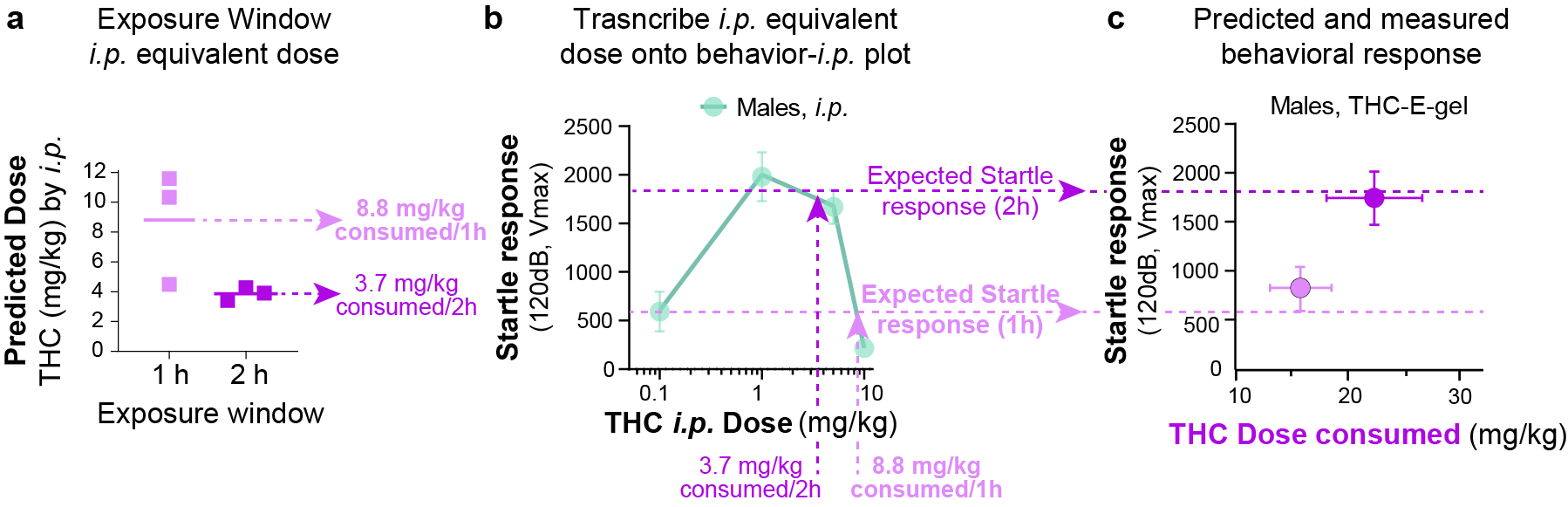

**Supplementary Figure S5: Methodology for THC-E-gel prediction of a behavioral response**

**a**) *i.p.* equivalent dose calculated from the triad of cannabimimetic behaviors in Figure 4. **b**) Transcribe *i.p.* equivalent doses to the THC-*i.p.* doe response plot for the given behavior to find the predicted behavior after 1 h or 2 h exposure to 10 mg/15 ml THC-E-gel. **c**) Predicted response (dashed line) and measured behavioral responses.

**Supplementary Table S6: Brain concentrations of THC and metabolites by sex**

**Supplementary Table S6: Brain tissue concentrations of THC and metabolites by sex**

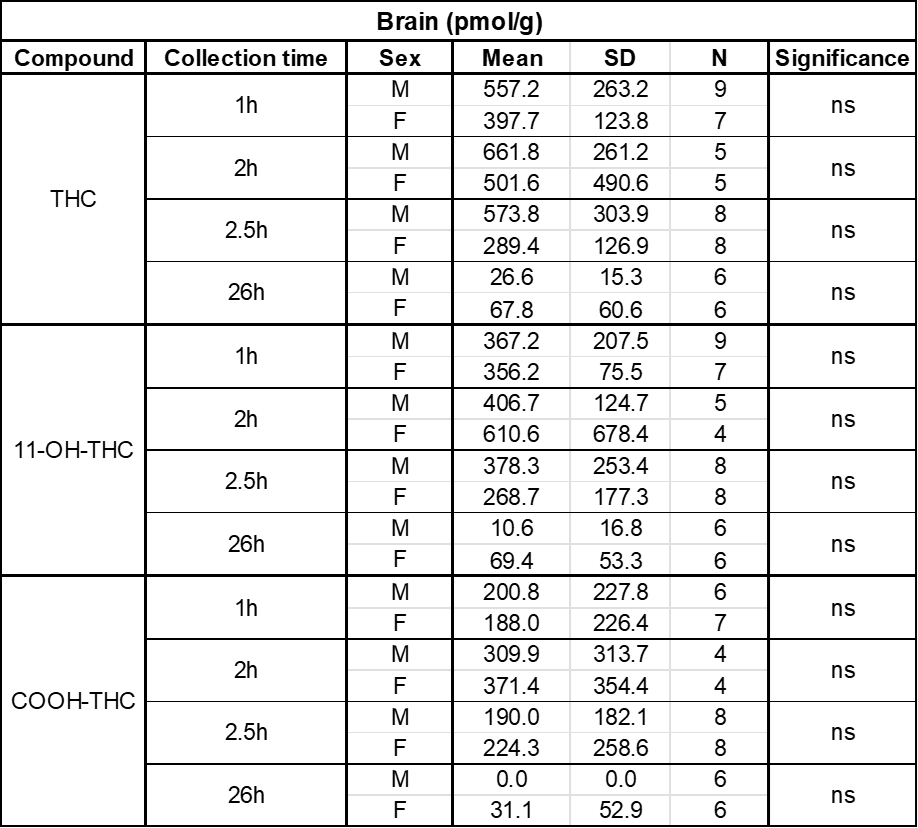

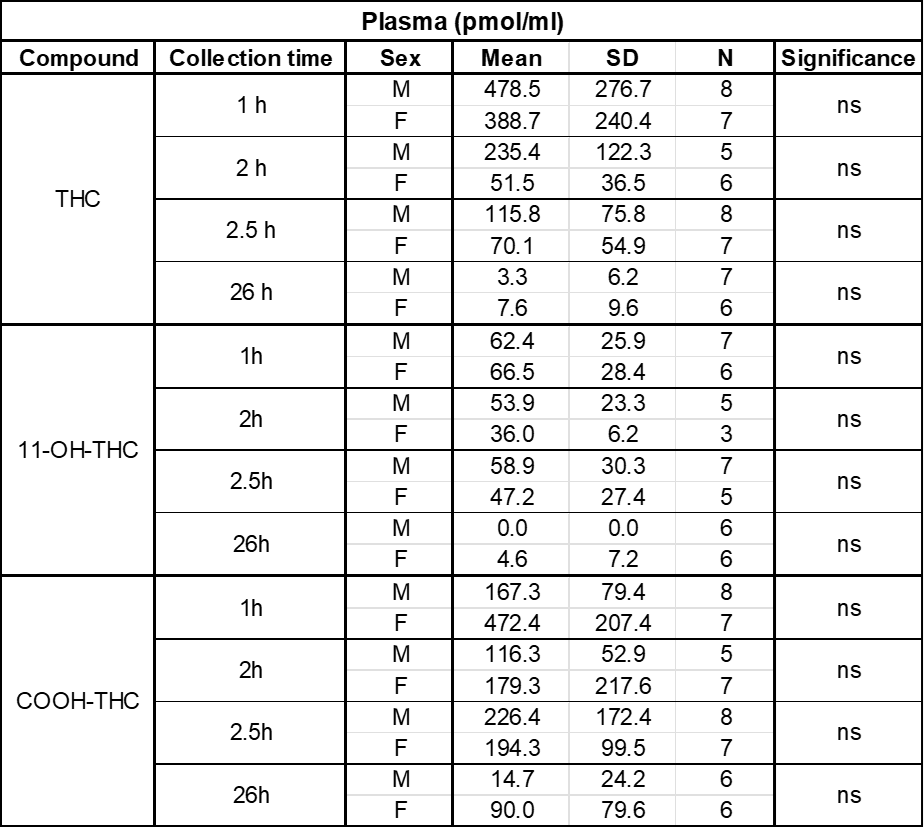

**Supplementary Table S7: Plasma concentrations of THC and metabolites by sex**
